# Yellow Fever Vaccine Protects Resistant and Susceptible Mice Against Zika Virus Infection

**DOI:** 10.1101/587444

**Authors:** Ana C. Vicente Santos, Francisca H. Guedes-da-Silva, Carlos H. Dumard, Vivian N. S. Ferreira, Igor P. S. da Costa, Ruana A. Machado, Fernanda G. Q. Barros-Aragão, Rômulo L. S. Neris, Júlio S. dos-Santos, Iranaia Assunção-Miranda, Claudia P. Figueiredo, André A. Dias, Andre M. O. Gomes, Herbert L. de Matos Guedes, Andrea C. Oliveira, Jerson L. Silva

## Abstract

Zika virus (ZIKV) emerged as an important infectious disease agent in Brazil in 2016. Infection usually leads to mild symptoms but severe congenital neurological disorders and Guillain-Barré syndrome have been reported following ZIKV exposure. The development of an effective vaccine against Zika virus is a public health priority, encouraging the preclinical and clinical studies of different vaccine strategies. Here, we describe the protective effect of an already licensed attenuated yellow fever vaccine (17DD) on type-I interferon receptor *knockout* mice (A129) and immunocompetent (BALB/c) mice infected with ZIKV. Yellow fever virus vaccination results in robust protection against ZIKV, with decreased mortality in the A129 mice, a reduction in the cerebral viral load in all mice, and weight loss prevention in the BALB/c mice. Despite the limitation of yellow fever (17DD) vaccine to elicit antibody production and neutralizing activity against ZIKV, we found that YF immunization prevented the development of neurological impairment induced by intracerebral virus inoculation in adult. Although we used two vaccine doses in our protocol, a single dose was protective, reducing the cerebral viral load. Different Zika virus vaccine models have been tested; however, our work shows that an efficient and certified vaccine, available for use for several decades, effectively protects mice against Zika virus infection. These findings open the possibility for using an available and inexpensive vaccine to a large-scale immunization in the event of a Zika virus outbreak.

## Introduction

Zika virus (ZIKV) probably emerged in the early 1900s and remained undetected for several years (1). This virus was first isolated in 1947 from a sentinel Rhesus monkey (*Macaca mulatta*) presenting febrile illness in Zika Forest, Uganda (2). The first case of ZIKV in humans was reported in 1952 (3), and was historically regarded as a self-limiting disease. However, the scenario began to change in 2013, when a large outbreak in French Polynesia was associated with cases of Guillain-Barré syndrome (4) and during the outbreak in Brazil (2014-2015), authorities reported an increased number of children born with microcephaly (1,5). Nowadays, infection by ZIKV is known to be associated with congenital neurological disorders (6).

ZIKV presents tropism for developing neurological cells and reaches the central nervous system (CNS) following infection (7, 8, 9, 10). ZIKV infection can lead to neuronal cell death and induce a proinflammatory state that affects the cell environment and consequently its development (8,10). The impairment in neuronal differentiation and proliferation induced by ZIKV infection can lead to a number of clinical consequences (e.g., microcephaly), which are known collectively as congenital Zika syndrome (8,10). Due to the devastating consequences of ZIKV infection, the WHO considers the development of preventive and therapeutic solutions a priority (9).

Different vaccine models, including inactivated and attenuated models, have been tested in preclinical studies (1, 11, 12). Some of these models have shown success in mice, and some of them have advanced to the clinical stage (1, 11, 12). It is known that infectious agents may lead to protection against other different but similar infectious agents (13). This mechanism is known as cross-protection and was, for example, the basis of the first vaccine developed, which led to the global eradication of smallpox (9). Members of the *Flaviviridae* family show similarity, and some members of this family are the targets of currently available vaccines, such as the attenuated yellow fever virus (YFV) vaccine (14). In the event that an already licensed vaccine effectively protects against ZIKV infection, several steps in the development of a new vaccine candidate could be skipped, and the vaccine already available on the market could be used. Northeastern Brazil was the region with the highest incidence of microcephaly between October 2015 and March 2015 and low coverage of YFV vaccination (15).

Here, we evaluated whether a vaccine for YFV, a flavivirus very similar to ZIKV, could prevent or at least decrease the severity of disease caused by ZIKV via a mechanism of cross-protection. We used the attenuated YFV 17DD vaccine because it is a vaccine model long used in humans with well-established tolerability. Vaccinated mice presented strong protection against ZIKV challenge, with lower mortality and cerebral viral loads. In addition, the mice were protected against neurological clinical signs of disease. Considering that we used a susceptible mouse (IFN-α/β receptor deficient, A129) and intracerebral inoculation, which causes a particularly serious illness, our results support the conclusion that YFV 17DD is a highly protective vaccine against Zika. This vaccine has the advantage of being already licensed and can be safety used in humans. These results indicate that we already have a vaccine against ZIKV infection and with an adequate program for vaccination and population awareness, we will be ready to combat a new ZIKV outbreak.

## Results

### YFV vaccine is safe for use in A129 mice and BALB/c mice

Based on a hypothesized cross-reaction between YFV vaccine and ZIKV, we evaluated the tolerability of the attenuated vaccine YFV 17DD in A129 mice, monitoring both weight loss and mortality after two immunization doses of the YFV vaccine. We tested three different doses of the YFV vaccine: 10^5^, 10^4^ and 10^3^ infectious viral particles (PFU). Only at the dose of 10^5^ PFU, although there was no difference in weight change (Figure 1A), we can observe 35% death (Figure 1B). In contrast, nonimmunized animals challenged with ZIKV lost weight (Figure 1A) and died (Figure 1B). The animals vaccinated with the 10^4^ and 10^3^ PFU doses of the YFV vaccine were asymptomatic. As 10^4^ PFU of the YFV vaccine did not induce apparent effects, we adopted this dose for subsequent experiments. We also evaluated the vaccine in BALB/c mice, and we did not observe any clinical signs of disease or death (data not shown).

**Figure 1:**
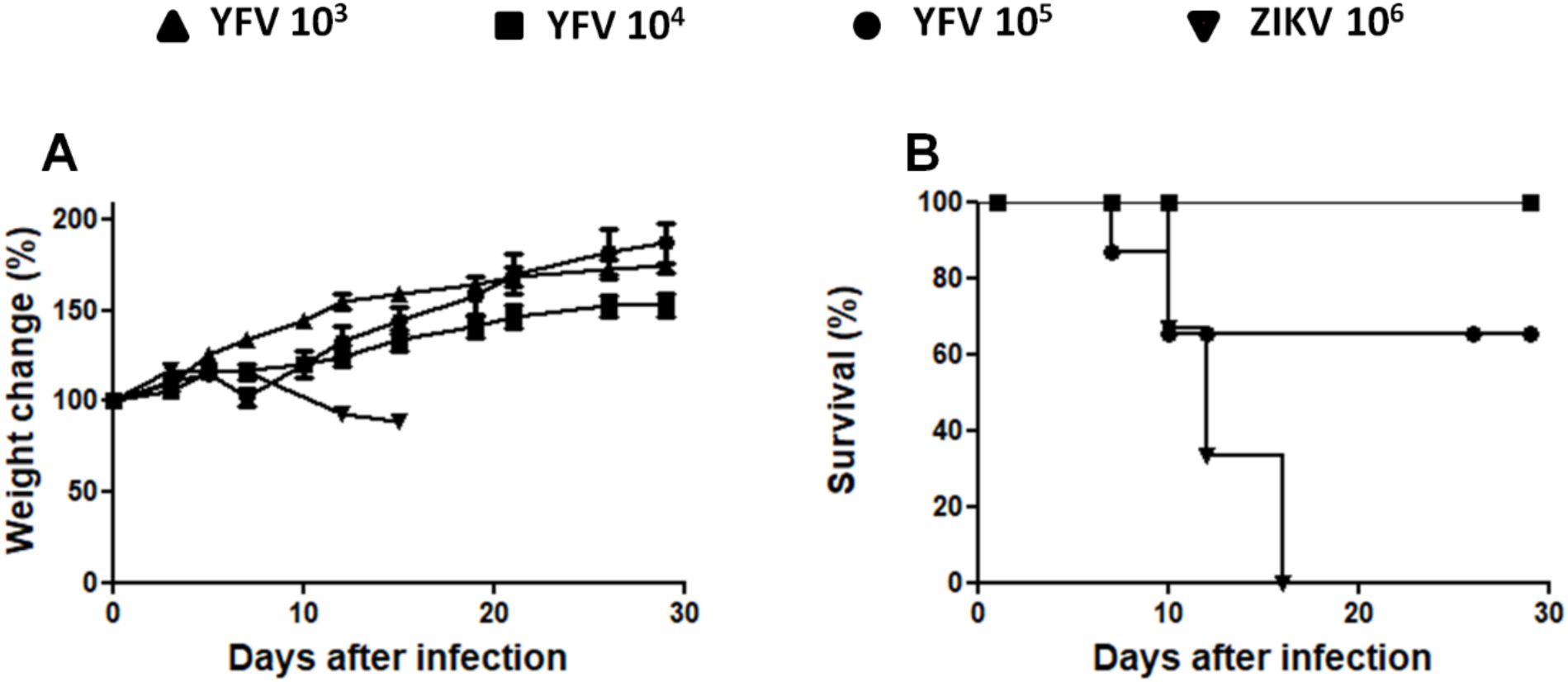
Dosing analysis of the YFV vaccine (subcutaneous route) in interferon-1 receptor knockout mice (A129). Mice were subcutaneously vaccinated with different doses of YFV 17DD (10^5^, 10^4^, or 10^3^ PFU) or challenged subcutaneously with ZIKV (10^6^ PFU). Weight **(A)** and survival **(B)** were measured. N= 5; Statistical Analysis: For weight change we used the One way ANOVA test, and no statistical difference was observed. For survival, the log-rank (Mantel-Cox) test was used. \*\**p*<0.01.

### YFV vaccine induces protection against ZIKV infection in A129 mice

The susceptibility of the A129 strain to ZIKV infection has been demonstrated previously (16) (Figure 1), making A129 mice a useful model to study ZIKV infection. We immunized A129 mice twice with the YFV vaccine or saline, which was used as a control. Seven days after the booster immunization, the mice were infected via the intracerebral route (IC) (as this route induces more rapid evolution and serious disease) (Figure 2). The attenuated YFV vaccine was shown to be effective in protecting the susceptible animals (Figure 3). The vaccinated mice group gained more weight (Figure 3A) and presented much lower mortality (Figure 3B) than the saline-treated mice group. The difference in mortality (Figure 3B) was more evident than the difference in weight loss (Figure 3A) since many of the unvaccinated mice lost weight rapidly and died within 10 days. Some of the mice that died after the tenth day lost less weight. However, the difference in weight loss was statistically significant.

**Figure 2:**
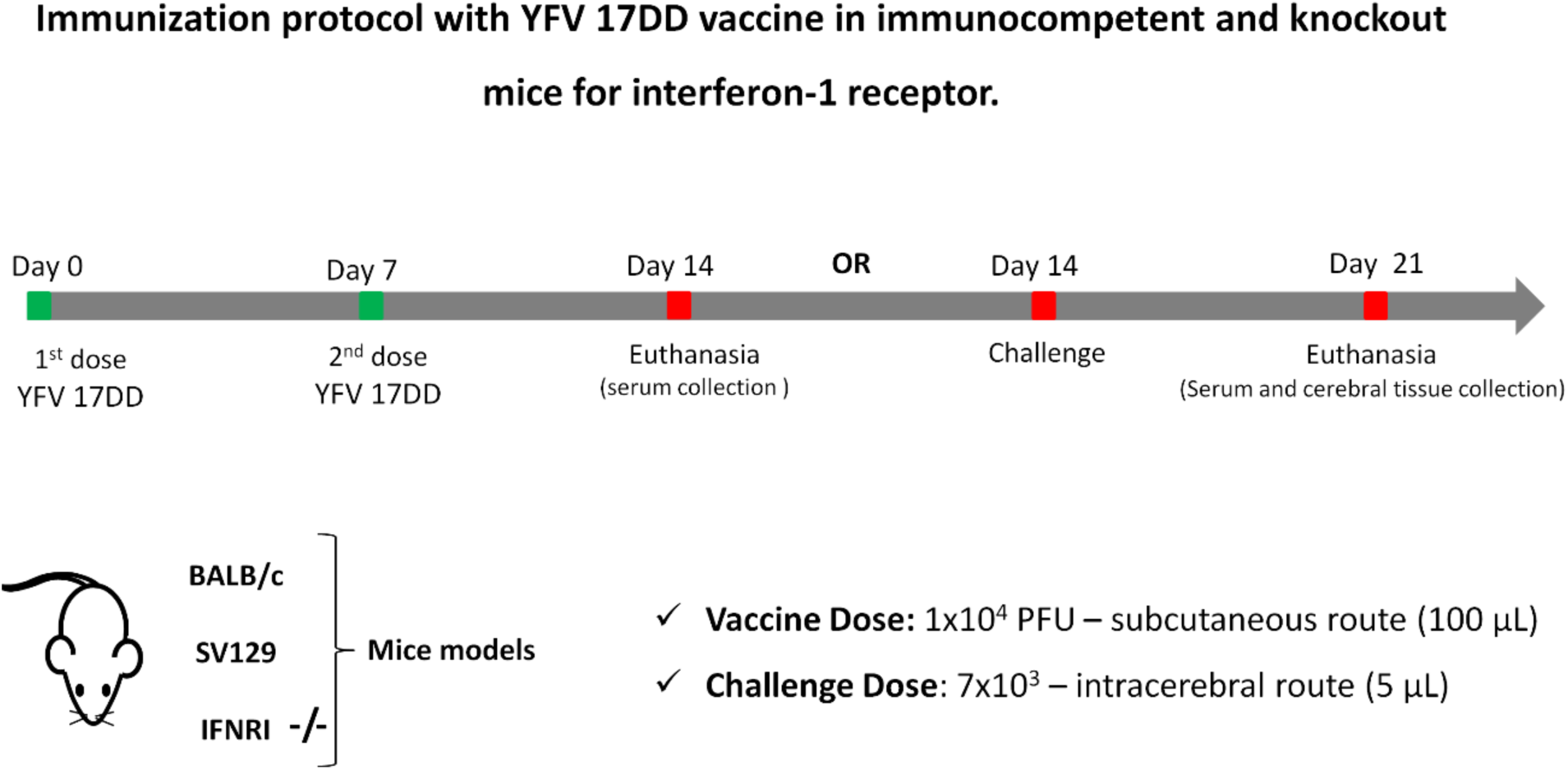
Immunization protocol with YFV 17DD in mice.

**Figure 3:**
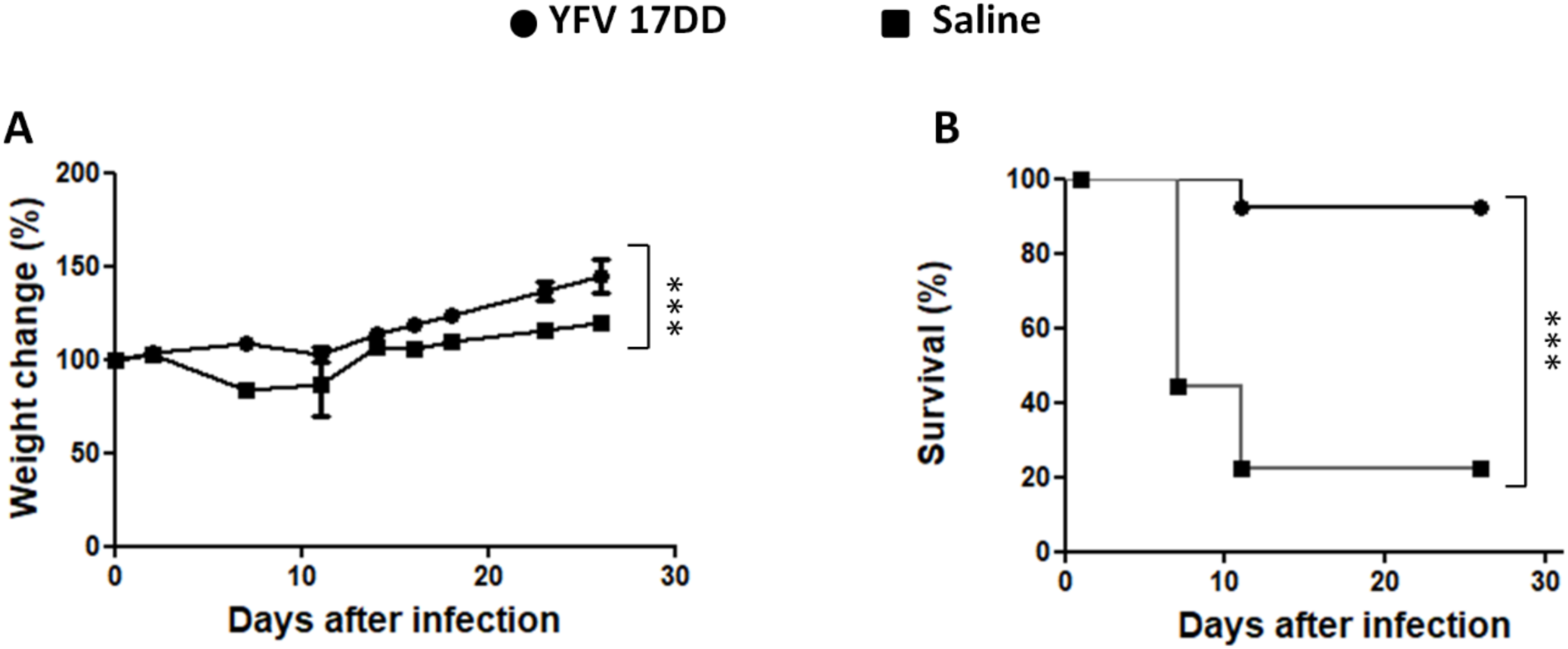
YFV vaccine protects interferon-1 receptor knockout mice (A129) against an intracerebral challenge with ZIKV. Mice were challenged via the intracerebral route with 7×10^3^ Zika virus particles. Weight **(A)** and survival **(B)** were measured. N=7; Statistical Analysis: Changes in weight were analyzed by two-way ANOVA, and survival was analyzed by the log-rank (Mantel-Cox) test. \*\*\**p*<0.0001.

### YFV vaccine induces protection against ZIKV infection in BALB/c mice

We also tested the YFV vaccine in immunocompetent BALB/c mice. The BALB/c mice were immunized twice and, after 7 days, challenged by the intracerebral route (following the same protocol used for the A129 mice as described on Figure 2). We observed that the vaccinated group presented no weight loss, while the saline group did (Figure 4A). The cerebral viral load was significantly different between the groups (Figure 4B), indicating that the prevention of clinical signs was correlated with lower viral propagation in the vaccinated mice.

**Figure 4:**
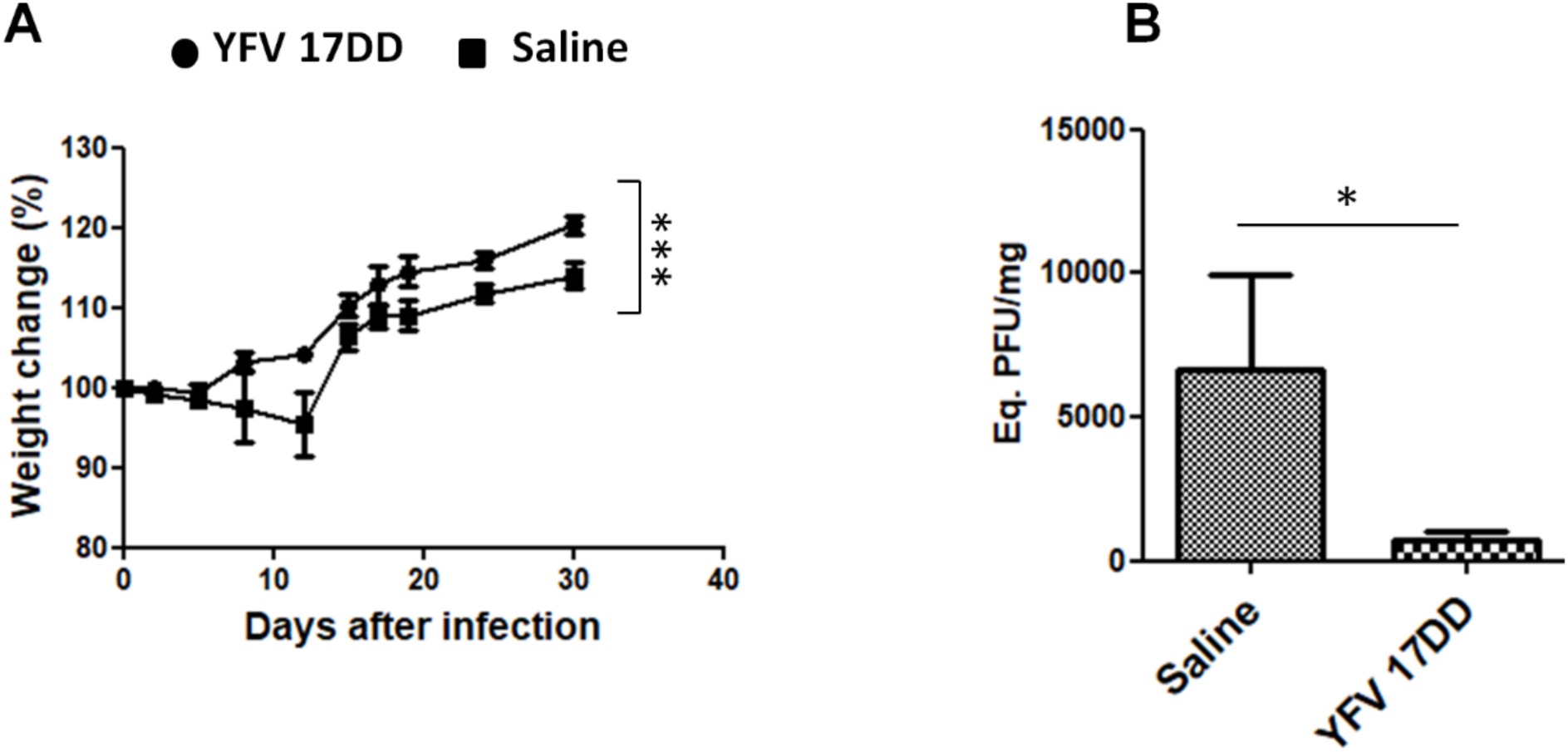
YFV vaccine protects immunocompetent BALB/c mice. Mice were challenged via the intracerebral route with 7×10^3^ Zika virus particles. Weight was measured **(A)**, and cerebral tissue qRT-PCR was performed 7 days after infection and ZIKV eq PFU/mg are shown **(B)**. N=5; Statistical Analysis: Changes in weight were analyzed by two-way ANOVA, and the qRT-PCR results were analyzed by the Mann Whitney test. ***p<0.0001, *p<0.05.

### YFV vaccine protects BALB/c mice against neurological signs

We observed different neurological disturbances, such as spinning when suspended by the tail, shaking, hunched posture, ruffled fur and paralysis, during ZIKV infection in the BALB/c mice. We evaluated these manifestations in the vaccinated and saline groups after challenge. All extremely recognizable clinical neurological signs were present in the saline group and completely absent in the vaccinated group (Table 1). In the saline group, 3 of the 5 animals presented an unsteady gait, marked by paralysis in at least one of the segments. In the vaccine group, no animals presented this clinical sign. In 2 of the 3 symptomatic mice, the unsteady gait was established as a permanent sequela (observed from 5 days after infection onwards). All mice in the saline group exhibited agitation and touch sensitivity, but all animals recovered from these behaviors. To assess motor behaviors, the animals were suspended by the tail, and 3 animals in the saline group showed the behavior of spinning during tail suspension. In 2 of these 3 animals, this behavior remained as sequelae (observed from 5 days after infection onwards). In the vaccine group, no mice exhibited this behavior. These results indicate that the mechanism of protection is efficient to control viral replication and brain damage, guaranteeing physiological homeostasis.

**Table 1.**
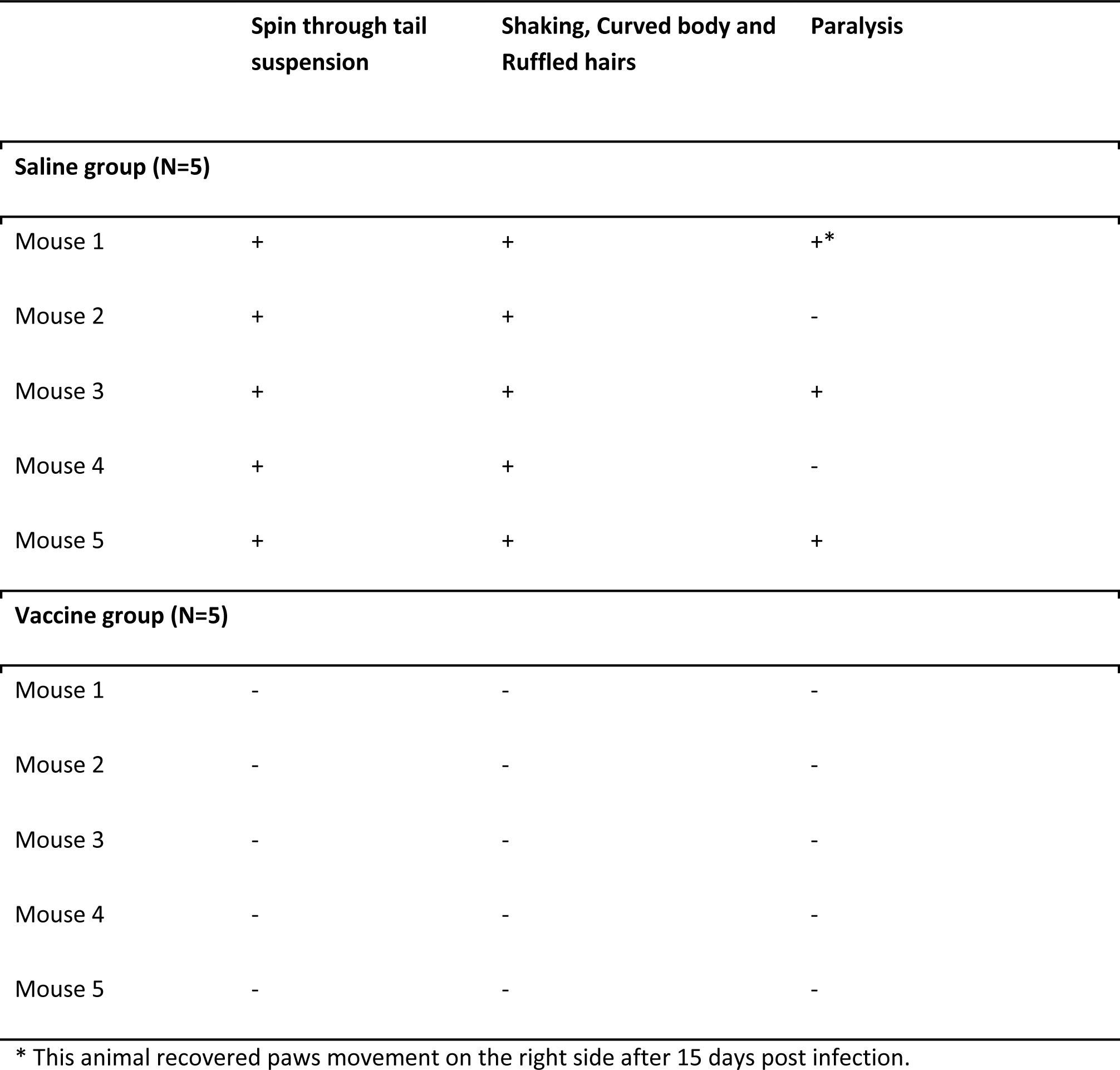
Neurological signs in immunocompetent BALB/c mice after infection. Mice were evaluated for the presence (+) or absence (-) of neurological signs, by two independent observers. Signs were evaluated daily from the first day after infection

### Low cross-reactivity based on antibody production and no microneutralization

We evaluated the capacity of the antibodies produced against YFV to cross-react with ZIKV. We observed that the immunization of BALB/c mice induced a small production of specific IgG antibodies against heterologous antigens (ZIKV) and homologous antigens (YFV) (Figure 5A and 5B), that could be detected 7 days after the booster immunization with significant differences between the experimental groups (Figure 5A and Figure 5B). This result indicates that the heterologous agent used in the vaccine (YFV) is able to elicit the production of a low level of antibodies that bind to ZIKV. We also evaluated the capacity of the antibodies produced against YFV to neutralize ZIKV infection in Vero cells. Our results demonstrated that the serum from the vaccinated mice did not neutralize ZIKV infection (Figure 5C), whereas the serum from the mice infected with ZIKV did, suggesting that the mechanisms induced by YFV can be related to the cellular immune response.

**Figure 5.**
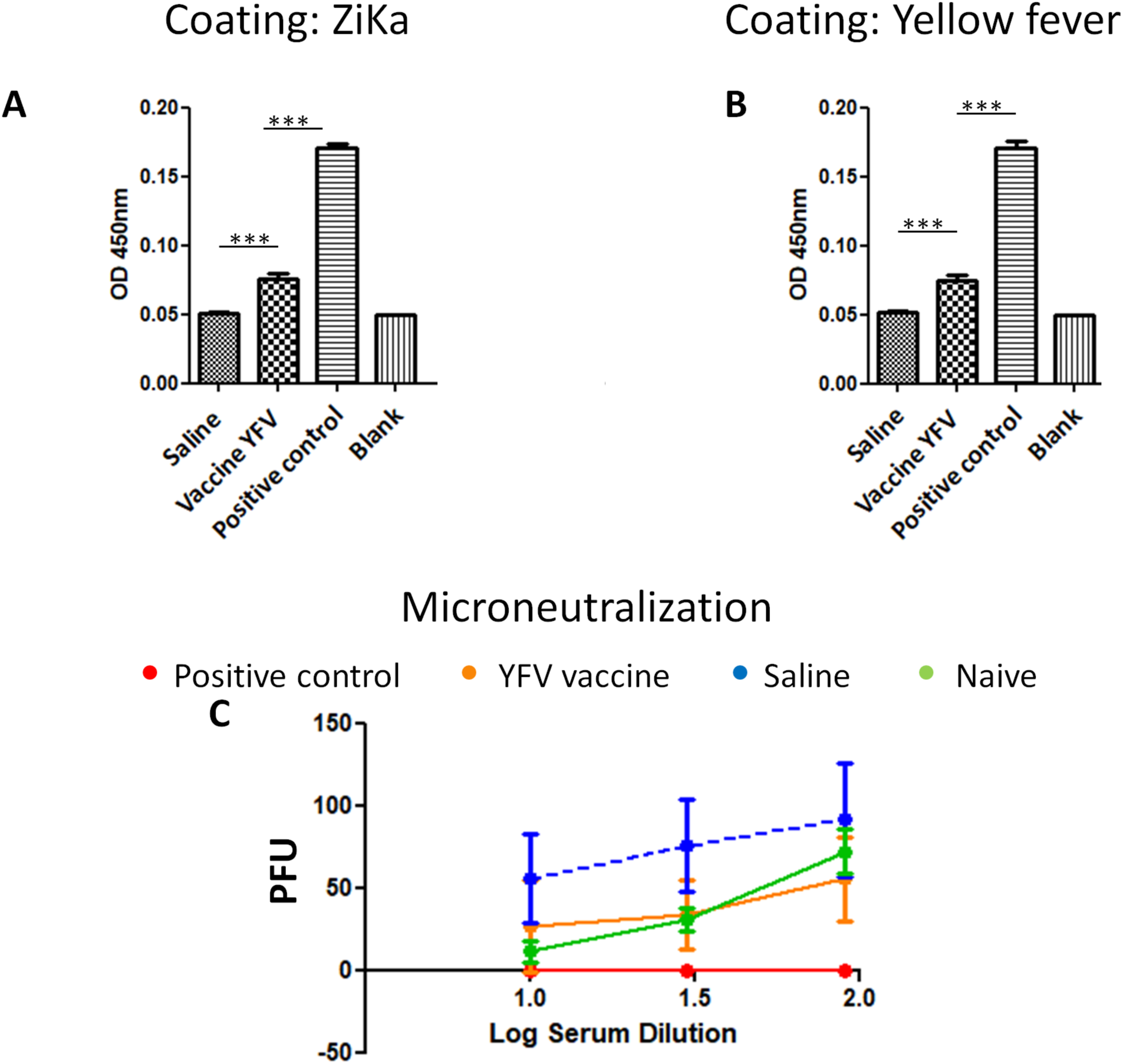
YFV vaccine elicit specific IgG response in immunocompetent mice. After the seventh day of the second dose, sera samples were collected and used to analyze antibody response at 1:360 dilution. (A) Antibody response by ELISA using ZIKV coating. (B) Antibody response by ELISA using virus YFV coating. All sera samples differed to saline group *** *p=*0.0001. One way ANOVA and Tukey post-test. (C) Sera were analyzed by its capacity to block zika infection by Microneutralization assay. PFU – plaque forming units (greater PFU indicate less capacity to block infection). Positive control was obtained by 4 consecutives infections in mice (separated by 10 days each), and collected 10 days after the fourth infection. For microneutralization all group differed from positive control. For both ELISA and microneutralization N=10.

### One YFV immunization is sufficient to induce protection

To evaluate whether a single dose of the YFV vaccine is enough to protect against ZIKV infection, SV129 (A129 background) (Figure 6A) and A129 (Figure 6B) mice were used. The mice were vaccinated, and, after 7 days, challenged by the intracerebral (IC) route with 7×10^3^ PFU ZIKV viral particles. Evaluation of the cerebral viral load performed demonstrated differences between the groups, with high viral loads in the saline groups in comparison to vaccinated group. When we evaluated A129 mice, we observed a significant reduction of viral load in vaccinated mice in comparison to saline group (Figure 6B). When we evaluated SV129 (WT mice), we observed a reduction of viral load without statistically difference (*p*=0.0556) (Figure 6A). These results suggest that one dose of the vaccine may be enough to confer protection.

**Figure 6.**
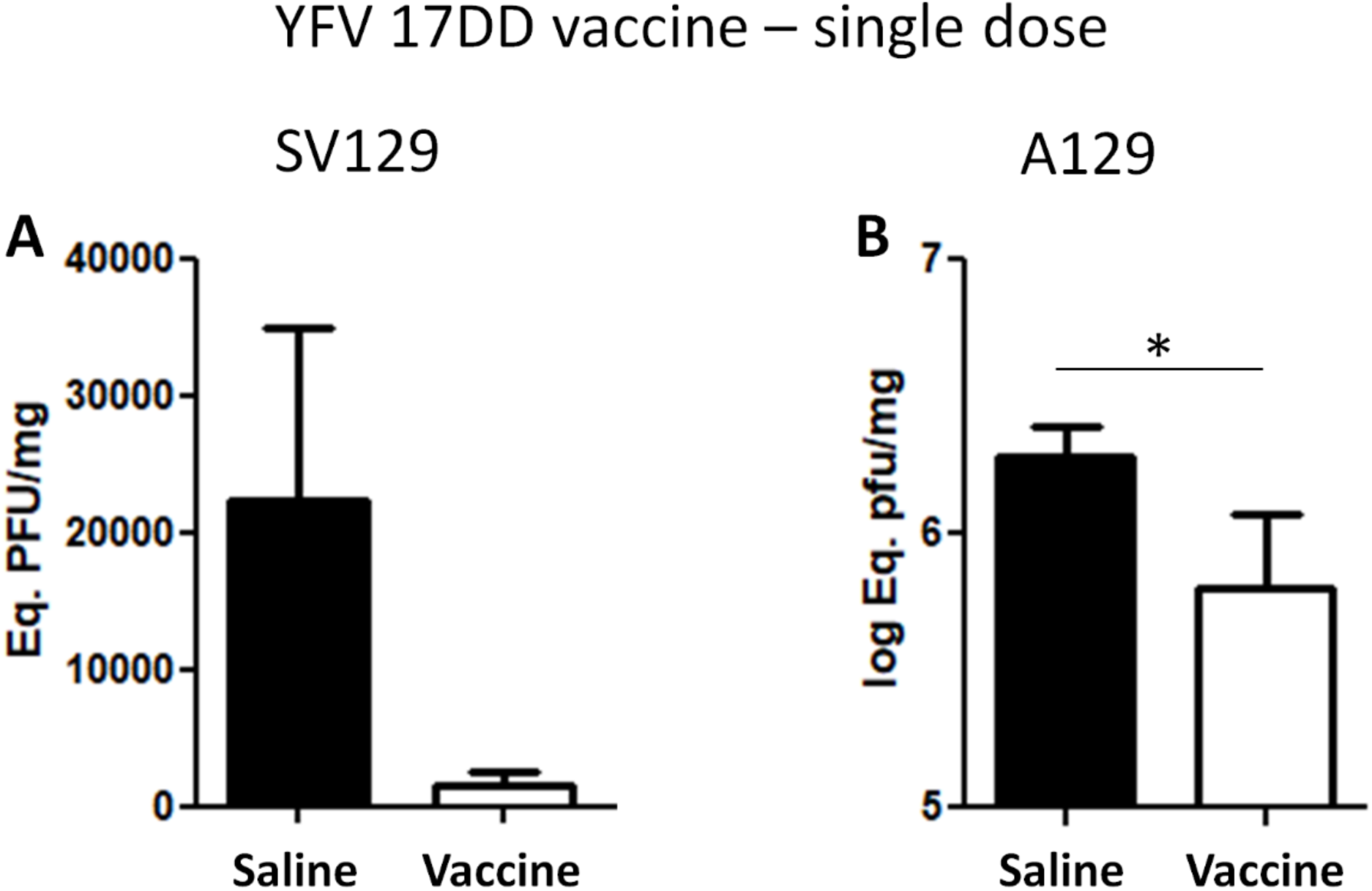
Immunization using a single dose of YFV in immunocompetent SV129 (A) and immunocompromised A129 (B) mice. Mice were immunized only one time and were challenged seven days after via the intracerebral route with 7×10^3^ ZIKV particles. Cerebral tissue qRT-PCR was performed 7 days after infection and the ZIKV eq PFU/mg are shown. N=5; Statistical Analysis: Mann-Whitney test. Although no significant difference was found for SV129 mice, the result was borderline significant (*p*=0.0556). \**p*<0.05.

## Discussion

We hypothesized that vaccination using YFV could protect against ZIKV infection through a cross-reaction mechanism. In this study, we immunized mice with the attenuated YFV 17DD vaccine and challenged them with ZIKV infection via the intracerebral route. IC infection, where ZIKV is inoculated directly into the central nervous system, is a highly invasive and pathogenic route; this route is considered the severe model of infection (17) and may require a strong immune response, which can probably only be achieved using live vaccines, to protect the brain. Attenuated agents, such as those in the polio and smallpox vaccines, have been used for years, including in mass campaigns, with great success, such as the global eradication of smallpox (9). YFV, for example, is one of the strongest immunogens ever developed, as it confers long-lasting protection with a single dose (14).

In our first step, we standardized the dose of YFV in A129 mice using immunization via a subcutaneous (SC) route. When we used 10^5^ PFU, YFV exhibited lethality in approximately 35% of the mice, but the animals that received 10^4^ or 10^3^ PFU were completely asymptomatic (Figure 1). Recently, a similar result was observed by another group using a chimeric attenuated vaccine (ChimeriVax-Zika) based on YFV with ZIKV epitopes (premembrane and envelope genes from YFV replaced by those from ZIKV) (18), which demonstrated that a dose of 10^5^ PFU resulted in a low mortality rate. This result is not a surprise since attenuated vaccines, despite being safe, require some precautions be taken for their use. When the tolerability of YFV (17-D) and ChimeriVax-Zika (CYZ) was analyzed in mice, CYZ was safer, inducing few deaths (18). However, the study comparing the two vaccines injected 5-day-old mice via the IC route to evaluate tolerability. Although this method of evaluation is important, it does not reflect the real world since vaccination does not occur via this route and is not performed in neonates. However, the YFV vaccine is recommended for people aged 9 months or older and has been used in pregnant women without any apparent adverse effects on fetuses (in this last group, vaccination may be discussed with the medical doctor). In addition, a number of other attenuated vaccines are used (e.g., polio, mumps, measles, rubella, BCG, and influenza), supporting the use of the YFV vaccine in humans and showing the safety of this vaccine approach.

We immunized A129 mice with 10^4^ PFU of YFV and challenged them with ZIKV via the IC route. Vaccination was shown to induce a high level of protection in the A129 mice. Giel-Moloney study using ChimeriVax-Zika showed a reduction in the viral load in vaccinated A129 mice; however, no survival results were reported (18). We demonstrated here that it is not necessary to use a chimeric vaccine because using YFV 17 DD is sufficient to induce protection against ZIKV. A study generated and evaluated a live attenuated vaccine candidate containing a 10-nucleotide deletion in the 3′ untranslated region (3’UTR) of the ZIKV genome (10-del ZIKV) as a vaccine against ZIKV. After immunization using 10-del ZIKV via the SC route and ZIKV challenge via the intraperitoneal (IP) route, the study authors demonstrated a strong reduction in the viral load; however, they did not report any survival studies (19). The protective efficacy of a live attenuated ZIKV vaccine with mutations in the NS1 gene and 3’UTR of the ZIKV genome was evaluated in only pregnant women, which did not allow us to compare those study results with our results (19,20). In our work, YFV provided protection in immunocompromised mice infected by the IC route, and this protection was demonstrated by a reduction in the viral load in the brain and by increased survival, as mortality was significantly lower in the vaccinated group (90% protection) than in the saline group, which is strong evidence of robust acquired immunity.

We also evaluated immunocompetent BALB/c mice. Recently, BALB/c mice were demonstrated to die after intracerebral infection using 10^3^ or 10^4^ PFU, and some of the group infected with 10^*2*^ PFU of ZIKV strain MR766 also died (Uganda, 1947) (21). Other immunocompetent mice also present mortality when infected at neonatal stage, such as Swiss (22). We observed that BALB/c mice did not die after IC challenge with ZIKV, and our model allowed us to study neurological disorders represented by easily recognizable clinical signs. The BALB/c mice immunized with YFV and challenged with ZIKV via the intracerebral route were effectively protected, exhibiting decreased weight loss and a reduced ZIKV cerebral load. Vaccination prevented the BALB/c mice from developing neurological disorders. Vaccination efficiently blocked viral propagation, which positively correlated with the clinical signs found in the BALB/c mice. Protection against IC challenge requires a potent immune response not only because this route causes more severe disease but also because the central nervous system presents some level of isolation from the rest of the body (immunoprivileged site).

Vaccines against ZIKV have been studied since the outbreak in 2015. Different approaches, including using a virus inactivated by formalin and subunit or DNA vaccines, have been tested (1, 11, 12, 23). Although of a different efficacy, an attenuated vaccine that induces a strong cellular immune response is desired. The attenuated YFV vaccine has been demonstrated to be effective in protecting against YFV using only one immunization dose (14). In a previous study, we used three doses of a pressure-inactivated vaccine with great success (24). Here, we evaluated one immunization protocol using immunocompetent (SV129) and immunocompromised mice (A129), and the ZIKV cerebral load was lower in the vaccinated SV129 and A129 mouse groups than in the corresponding saline groups (but the difference was statistically significant only between the A129 mouse groups). This result indicates that one dose is sufficient to generate an immune response that decreases cerebral viral propagation.

The mechanism of YFV vaccination that protects against YFV infection also involves neutralizing antibodies (25). CYZ has been shown to elicit antibodies in mice and reduce the viral load in a vaccinated group (18). When we evaluated antibody production against ZIKV, we detected little production, and the antibodies did not have the capacity to neutralize ZIKV infection in Vero cells. It is important to note that the vaccination protocol used did not favor antibody production because the time between immunization and challenge was too short (i.e., we also did not observe a high amount of antibodies against YFV). These results indicate that the mechanisms of protection do not involve antibodies.

We suggest that the mechanism of protection is associated with the cellular response. The YFV vaccine YF-17D induces a robust cellular immune response through the activation of a mixed Th1 and Th2 response, cytotoxic CD8+ T cells and a neutralizing antibody response (26). These mixed responses are elicited by the activation of toll-like receptors (TLRs) such as TLR2, TLR3, TLR7, TLR8 and TLR9 on dendritic cells (27). A study using antigen-specific, interferon-γ (IFN-γ)-secreting MYD88 -/- CD4+ T cells and CD8+ T cells (27) indicated that innate immunity has the role of inducing adaptive immunity during infection by attenuated YFV. Several CD4+ and CD8+ T cell epitopes have been characterized and related to the protection induced by YFV vaccines (28, 29). Recently, the importance of the cellular response against ZIKV infection has been assessed, and CD4+ T cells (30,31,32), CD8+ T cells (32) and several epitopes involved in the response have been characterized (31). Based on the absence of neutralizing antibodies against ZIKV after YFV vaccination, we suggest a cross-reaction involving CD4+ and CD8+ T cells. Cross-reactivity based on T cells has already been demonstrated (33).

Many ZIKV vaccine candidates are in the preclinical phase, and some are in clinical phases I and II. Different technologies, such as live attenuated vaccines, recombinant vector vaccines, subunit vaccines, whole inactivated vaccines, mRNA vaccines and DNA vaccines, have been tested (1, 11, 12, 23). Undoubtedly, the study and development of new vaccines are extremely important, as these processes allow us to have more efficient and safer models. Some of these models may turn out to be highly effective vaccines, and some may not, but it will still take time to make these vaccines available. This gap can be filled by the YFV vaccine, which has been successfully used for decades in the human population and is readily available now. It is possible that the YFV vaccine may be effective in protecting humans against ZIKV, especially against neurological diseases in adults and congenital Zika syndrome. A recent epidemiological study reports that preexisting infection with dengue virus (as determined by high antibody titers) was associated with reduced risk of ZIKV infection (34). Also, based on epidemiological data, it has been suggested that pregnant women in regions of Brazil with lower YFV vaccination coverage are at higher risk for the development of microcephaly (15). However, no experimental evidence has been provided for this hypothesis. If YFV really protects against ZIKV in humans, it would be an immense advantage for the YFV vaccination model since its pros and cons in clinical practice are already well known. In addition, the YFV vaccine would be a vaccine capable of protecting against two distinct pathogens simultaneously. Substantial time and resource savings could be accrued by using an already licensed vaccine.

## Conclusion

YFV vaccination is protective against ZIKV infection in resistant and susceptible mouse models that underwent one or two immunizations.

## Materials and methods

### Cells

Vero (African green monkey kidney) cells (CCL 81) were obtained from the American Type Culture Collection (ATCC), Manassas, VA, EUA, and cultured in high-glucose Dulbecco’s modified Eagle’s medium (Gibco™ DMEM; Thermo Fisher Scientific - Manassas, VA, USA) The culture medium was supplemented with 10% fetal bovine serum (FBS; Vitrocell Embriolife, Campinas, SP, Brazil) and 100 µg/mL streptomycin, and the cells were maintained at 37°C in a 5% CO_2_ atmosphere.

### Mice

We used different mouse strains in this study: the immunocompetent BALB/c and SV129 strains and the immunocompromised A129 strain (IFNAR1). All animals were obtained from the UFRJ’s Central Biotherm (Rio de Janeiro/RJ, Brazil). All procedures were performed according to the guidelines established by the Ethics Committee for Animal Use of UFRJ (CEUA 069 /16).

### ZIKV and YFV

The strain of ZIKV used in this study was ZIKVPE_243_ (Genbank ref. number KX197192) isolated from a febrile case in the state of Pernambuco, Brazil, and was kindly given by Dr. Ernesto T.A. Marques Jr. (Centro de Pesquisas Aggeu Magalhães, FIOCRUZ, PE, Brazil), and of YFV was YFV 17DD, which was kindly given by LATEV, Bio-Manguinhos/Fundação Oswaldo Cruz (Rio de Janeiro/RJ, Brazil). The viruses were propagated as described previously (22,24), and viral titers were determined in Vero cells using a standard plaque assay at day 5 post-infection with crystal violet staining (Merck Millipore). The viral titers were determined in aliquots of harvested medium, and stocks of the viruses were stored at −80°C.

### Safety study

For the safety study, we injected the YFV vaccine at the doses 10^3^, 10^4^ and 10^5^ PFU via the SC route into A129 mice, and we challenged the mice with ZIKV as a control.

### Vaccination and challenge

We performed two immunizations with attenuated YFV via the subcutaneous (SC) route using a dose of 10^4^ PFU with 7-day intervals between the doses. Mice were challenged with ZIKV by inoculating 5 μL of ZIKV (7×10^3^ viral particles) via the intracerebral (IC) route using a 0.5 mL Hamilton syringe and 27 G ¼ needles. Control mice were treated with phosphate-buffered saline (PBS) instead of YFV. The challenged mice were observed for 4 weeks to evaluate clinical signs, including ruffled fur, vocalization, shaking, hunched posture, spinning during tail suspension, paralysis and death. Dying animals were euthanized humanely. Protocol is summarized on Figure 2. Using an alternative protocol, mice received only one dose and seven days after were challenged.

### Determination of the viral load by qRT-PCR

Seven days post-immunization (booster) with the attenuated YFV vaccine, animals were challenged by the IC route. The viral load was measured in the brain tissue of the mice at day 7 post-challenge (peak viremia in these models) by qRT-PCR using primers/probes specific for the ZIKV E gene as previously described (22). Cycle threshold (Ct) values were used to calculate the equivalence of log PFU/mg tissue after conversion using a standard-curve with serial 10-fold dilutions of ZIKV stock sample.

### Enzyme-linked immunosorbent assay (ELISA) evaluation of anti-mouse IgG levels in the serum of immunized immunocompetent mice

Polystyrene microplates (Corning, New York, NY, EUA) were coated overnight at 4°C with 10^5^ ZIKV or YFV viral particles. After blocking for 2 h with PBS containing 1% bovine serum albumin (BSA) (LGC Biotecnologia, Cotia, SP), the serum from mice vaccinated with YFV were adsorbed in the wells at different concentrations and incubated overnight at 4°C. Then, peroxidase-conjugated goat anti-mouse IgG antiserum (1:4,000; Southern Biotech) was added to the wells, and the plate was incubated for an additional period of 1 h. Peroxidase activity was revealed via hydrogen peroxide and tetramethylbenzidine (TMB). The reaction was stopped with H_2_SO_4_ (2.5 N), and the optical density (OD) at 450 nm was determined with a spectrophotometer using SOFTmax PRO 4.0 software (Life Sciences Edition; Molecular Devices Corporation, Sunnyvale, CA).

### Microneutralization

In the microneutralization assay, serum samples were initially diluted 1:10 and then serially diluted in 2-fold steps. Then, the dilutions were mixed at a 1:1 volume ratio with approximately 150 PFU of ZIKV, and the samples were incubated for 30 min at 37°C. Then, the samples were incubated with 60-70% confluent Vero cells in 24-well culture plates for 1 h at 37°C and 5% CO_2_. Next, each well received 1 mL of high-glucose DMEM containing 1% FBS, 1% 100 μg/mL penicillin, 100 μg/mL streptomycin mixed solution (LGC Biotecnologia, Cotia, SP) and 1.5% carboxymethylcellulose (CMC; Sigma-Aldrich Co, Missouri, EUA). The plates were incubated at 37°C and 5% CO_2_ for 4 days. The cells were fixed by adding 1 mL of 4% formaldehyde for 30 min. Each plate was washed and stained with a crystal violet solution (1% crystal violet, 20% ethanol). The number of plaques in each well was counted to determine the neutralizing effect of the serum on ZIKV.

### Experimental procedure of the clinical analysis

After intracerebral challenge with ZIKV, animals were observed daily and analyzed for 60 min for clinical signs of infection by comparing the vaccinated infected and control groups with healthy mice. We qualitatively analyzed behavioral signs such as exploratory activity, vocalization, prostration, and alterations in the coat and the presence of a motor deficit, which is widely visible and associated with weight loss. The animals underwent tail suspension for a maximum of 60 seconds for the evaluation of neurological alterations. For this examination, the animals were tested twice daily with a minimum interval of 5 min between analyses. The temporal qualitative analysis of the clinical parameters showed that the vaccinated mice appeared more active than the nonvaccinated mice and were free of signs of disease in all evaluated parameters in the period of acute infection. After resolution of the acute infection, some animals in the control group continued to exhibit evident motor sequelae.

### Statistical analysis

Statistical analysis was performed using GraphPad Prism 6.01 (GraphPad). Data are reported as the mean ± SEM. Tests used: log-rank (Mantel-Cox), One way ANOVA with Tukey post-test, Two way ANOVA, and Mann Whitney.

### Financial support

This work was supported by grants from Conselho Nacional de Desenvolvimento Científico e Tecnológico (CNPq), Fundação Carlos Chagas Filho de Amparo à Pesquisa do Estado do Rio de Janeiro (FAPERJ), Ministério da Saúde (Decit/SCTIE/MS), Coordenação de Aperfeiçoamento de Pessoal de Nível Superior (CAPES) and Financiadora de Estudos e Projetos (FINEP) of Brazil.

## Acknowledgments

The authors thank Natália Cristina Cerne Barreto and Gileno dos Santos de Sousa for their competent technical assistance.

